# Rift Valley fever virus induces fetal demise through direct placental infection

**DOI:** 10.1101/383745

**Authors:** Cynthia M. McMillen, Nitin Arora, Devin A. Boyles, Joseph R. Albe, Michael R. Kujawa, Jeffrey F. Bonadio, Carolyn B. Coyne, Amy L. Hartman

**Affiliations:** Center for Vaccine Research, University of Pittsburgh, Pittsburgh, PA, USA; Department of Infectious Diseases and Microbiology, University of Pittsburgh School of Public Health, Pittsburgh, PA, USA; Department of Pediatrics, University of Alabama at Birmingham, Birmingham, AL, USA; Department of Pediatrics, University of Pittsburgh School of Medicine, Pittsburgh, PA, USA; Center for Microbial Pathogenesis, Children’s Hospital of Pittsburgh of the University of Pittsburgh Medical Center, Pittsburgh, PA, USA; Department of Pathology, Magee Women’s Hospital of the University of Pittsburgh Medical Center, Pittsburgh, PA, USA

## Abstract

Rift Valley fever virus (RVFV) infections in pregnant livestock are associated with high rates of fetal demise and have been linked to miscarriage in pregnant women. To address how acute RVFV infection during pregnancy causes detrimental effects on the fetus, we developed an immunocompetent pregnant rodent model of RVFV infection. We found that pregnant rats were more susceptible to RVFV-induced death than their non-pregnant counterparts and that RVFV infection resulted in intrauterine fetal death and severe congenital abnormalities, even in pups from infected asymptomatic pregnant rats. Virus distribution in infected dams was widespread, with a previously unrecognized preference for infection, replication, and tissue damage in the placenta. In human mid-gestation placental tissue, RVFV directly infected placental chorionic villi, with replication detected in the outermost syncytial layer. Our work identifies direct placental infection by RVFV as a mechanism for vertical transmission and points to the teratogenic potential of this virus in humans. This is the first time vertical transmission of RVFV has been shown in species other than livestock. This study highlights the potential impact of a future epidemic of this emerging mosquito-borne virus.

## Introduction

Rift Valley fever (RVF) is a veterinary disease of domesticated livestock that is frequently transmitted to humans. RVF virus (RVFV; Phenuiviridae; formerly Bunyaviridae) is currently endemic in many regions of Africa and is transmitted by a range of mosquito species (1-4). Prominent instances of emergence of RVFV in new areas (such as Egypt in 1977-78 and Saudi Arabia in 2000-01) have caused concern for further spread given that mosquito species found in Europe and the Americas could potentially harbor and transmit RVFV (5-7). The World Health Organization warns of a pending public health emergency caused by RVF due to insufficient vaccines and therapeutics (8).

Epizootic outbreaks of RVF predominantly affect young ruminants such as sheep, goats, cattle, and camels. Within a given outbreak, infected animals succumb to disease characterized by fulminant hepatic necrosis (9). The most notable and economically devastating outcome of RVF is the “abortion storm” that sweeps through herds of pregnant livestock, with abortogenic rates reaching as high as 90-100% in pregnant ewes (10,11). Even live-attenuated veterinary vaccines induce abortions in pregnant animals and cause physical abnormalities in fetuses and newborns (12).

In humans, RVF disease is primarily an acute febrile illness accompanied by body aches and joint pain, with occasional progression to more severe systemic disease. Data from human outbreaks, while limited, suggest that vertical transmission of RVFV to the developing human fetus can occur with detrimental outcomes. Two published cases of vertical transmission resulted in infection and pathological outcomes in the fetuses. In one instance, a pregnant woman became acutely infected with RVFV, resulting in delivery of an infant with rash, enlarged liver and spleen, and jaundice (13). In another family where 9 members recently contracted RVF (one of whom died), a pregnant woman displayed clinical signs of RVF a few days before labor and delivered an infant who subsequently died of RVF within a week (14). In a recent study in Sudan, pregnant women with confirmed RVF illness during pregnancy had higher rates of 2^nd^ and 3^rd^ trimester miscarriages or stillbirths (odds ratio of 7.4) (15).

Given the high incidence of fetal abortions in livestock and the potential for miscarriage in pregnant women, this study addressed vertical transmission of RVFV in non-livestock species and the susceptibility of human placental tissue to RVFV infection. Using immunocompetent Sprague-Dawley rats infected with a wild-type pathogenic strain of RVFV, we demonstrate direct vertical transmission and intrauterine fetal death similar to that observed in livestock. Importantly, vertical transmission occurred in pregnant dams with no clinical signs of disease, even after infection during late-gestation when the placenta is fully-formed. Antenatal infection resulted in delivery of stillborn pups with stunted development and gross anatomical changes. Remarkably, for a hepatotropic virus, the placenta had a higher viral burden than the liver and other maternal organs. *Ex vivo* inoculation of second-trimester human fetal tissue explants with RVFV resulted in active replication in the syncytiotrophoblast layer of the placenta, a structure typically resistant to viral infections. This is the first *in vivo* study to recapitulate the teratogenic effects of RVFV infection in livestock and the first time RVFV has been shown to directly infect human placental tissue. This study also highlights the previously unrecognized and potentially severe effects of RVFV infection in pregnant women.

## Methods

### Ethics Statement

All animal work described here was carried out in strict accordance with the Guide for the Care and Use of Laboratory Animals of the National Institutes of Health (NIH) and the Animal Welfare Act (AWA). The protocol was approved and overseen by the University of Pittsburgh Institutional Animal Care and Use Committee (IACUC). The Association for Assessment and Accreditation of Laboratory Animal Care (AAALAC) has fully accredited the University of Pittsburgh.

### Biosafety information

All work with live RVFV was conducted at BSL-3 in the University of Pittsburgh Regional Biocontainment Laboratory (RBL). For respiratory protection, personnel wore powered air purifying respirators (PAPRs; Versaflo TR-300, 3M) and used a class III biological safety cabinet. All animals were housed in individually-ventilated micro-isolator caging (Allentown, Inc). Vesphene IIse (1:128 dilution, Steris Corporation) was used to disinfect all liquid wastes and surfaces at risk of contact with the infectious agent. The RBL is a shower-out facility that requires a full clothing change into scrubs prior to entry and a personal shower and new scrubs upon exit. All solid wastes, used caging, and animal wastes, were steam-sterilized. Animal carcasses were incinerated or digested via alkaline hydrolysis (Peerless Waste Solutions). All tissues or samples destined for removal from BSL-3 were inactivated using methods described below; all inactivation methods have been verified and approved by a University of Pittsburgh biosafety oversight committee. The University of Pittsburgh Regional Biocontainment Laboratory is a registered entity with the Centers for Disease Control and Prevention and the United States Department of Agriculture for work with RVFV.

### Virus and cell culture

Virulent RVFV strain ZH501, generated from reverse genetics plasmids (16), was generously provided by Barry Miller (CDC, Ft. Collins, CO) and Stuart Nichol (CDC, Atlanta). Virus was propagated on Vero E6 (ATCC; CRL-1586) cells using standard methods. Viral titer was determined by standard viral plaque assay (VPA). Briefly, 200 μL tissue homogenates or supernatant serial-diluted in D2 media (Dulbecco’s Modified Eagle Medium, 2% FBS, 1% penicillin/streptomycin) was plated on 6-well plates (Corning) containing Vero E6 cells at 90-95% confluency. Plates were incubated (37°C, 5% CO_2_) for 1 hour for viral adsorption and rocked every 15 minutes. Following incubation, the virus inoculum was removed and replaced with 3 mL of agarose overlay (Minimum Essential Medium, 2% FBS, 1% penicillin/streptomycin, 1.5% (1M) HEPES buffer, and 0.8% SeaKem agarose). Plates were incubated for 3 days at 37°C, 5% CO2 to allow plaque formation. Plates were treated with 2 mL 10% formaldehyde for 4 hours for cell fixation and virus inactivation. Plaques were visualized after crystal violet (CV) staining (0.1% CV solution in 20% EtOH) and quantitated using the following equation: Average # plaques x 5 (dilution factor) x dilution = PFU/ml.

### Animals

Age-matched non-pregnant and time-mated Sprague Dawley rats (SD; 6-8 weeks) were obtained from Envigo Laboratories. A positive copulation plug verified pregnancy for early-gestation (embryonic day 5; E5) and late-gestation (embryonic day 14; E14) females. All pregnant rats were delivered to individual cages and non-pregnant rats were housed three to a cage in temperature-controlled rooms with a 12-hour day/12-hour night light schedule. Food (IsoPro Rodent 3000) and water were provided ad libitum. Rats were implanted with programmable temperature transponders (IPTT-300; Bio Medic Data Systems) subcutaneously between the shoulder blades. For infection, all rats were anesthetized by inhalation of isoflurane vapors (IsoThesia, Henry Schein) and inoculated subcutaneously in the hind flank with 500 μL of RVFV diluted in D2 media. Weight and body temperature were recorded daily starting the day of infection. Additionally, each animal was closely monitored twice daily for the development of clinical signs. Endpoint criteria, which prompts immediate euthanasia, was defined based on weight, temperature, appearance, and behavioral scoring parameters. Once euthanasia criteria were met, rats were anesthetized by inhalation of isoflurane vapors followed by an immediate blood draw and euthanasia by cardiac puncture.

Rats were inoculated on E14 with the following doses of RVFV strain ZH501: 7.5 x 10^1^ pfu (n=5), 1.8 x 10^2^ pfu (n=6), 1.5 x 10^3^ pfu (n=11) and 2.6 x 10^4^ pfu (n=6). Unless dams met euthanasia criteria, they progressed to full-term and delivered pups on E22 (8 dpi). After delivery, dams and pups were not disturbed until 5 days post-delivery (13 dpi) to reduce stress. Added stress on the dams can lead to consumption of newborn pups by dam (17, 18). Weight monitoring of pups began on neonatal day 5 (13 dpi) until 18 dpi when both dam and pups were euthanized at the predetermined end of the study (Fig. 1A).

**Figure 1:**
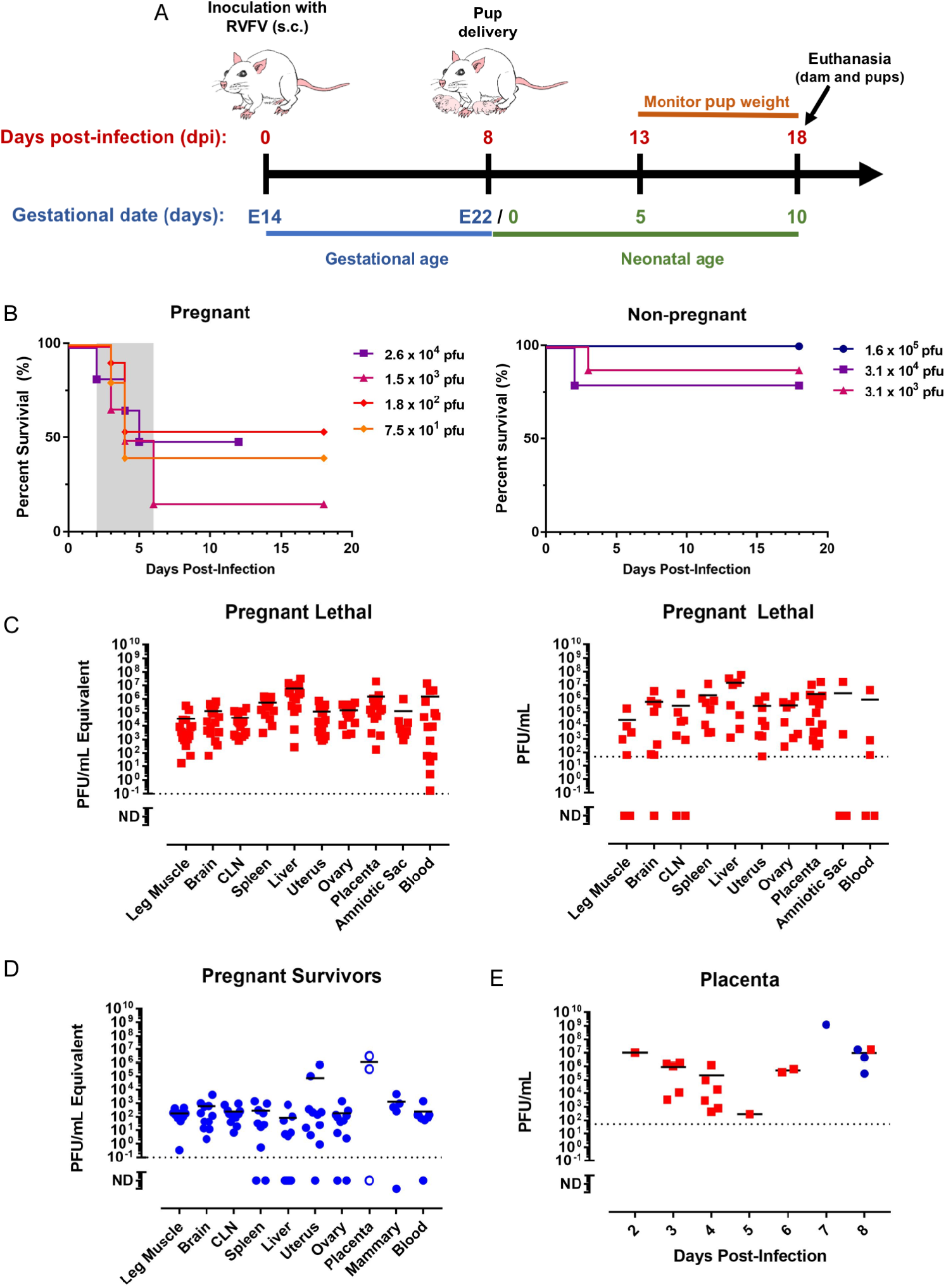
Pregnant rats are more susceptible to death after RVFV infection, with virus homing to the liver and placenta. (A) Experimental design for E14 SD rats infected with RVFV After delivery at E22, dams and pups were not disturbed until 5 days post-delivery (13 dpi). Euthanasia of surviving dams and pups occurred 18 dpi (10 days post-delivery). (B) Survival of RVFV-infected pregnant dams and non-pregnant SD rats (n=3-6 per dose). The shaded area represents the 2- to 6-day clinical window when lethally-infected pregnant rats were euthanized due to severe disease. (C) vRNA (q-RT-PCR; left) and infectious virus (VPA; right) in tissues from pregnant rats that succumbed (red squares; n=17) between 2-6 dpi. (D) vRNA in tissue samples from pregnant rats that survived infection (blue circles; n=11) and were euthanized 18 dpi. Placenta samples (open blue circles) were obtained at day of delivery (8 dpi). (E) Infectious virus measured by VPA in placental samples obtained from lethally infected (red squares) and surviving (blue circles) rats at the indicated day after infection. CLN; cervical lymph node. Dashed horizontal lines represent the limits of detection (LOD) of the q-RT-PCR (0.1 PFU equivalent/mL) and VPA (50 PFU). ND, not detected (below the LOD).

One rat at embryonic day 5 (E5) was infected with 1.5 x 10^5^ pfu of RVFV. A non-infected dam at E5 gestation was observed in parallel. Dams were euthanized at a pre-determined date of 7 days post-infection, corresponding to embryonic day 13.

Age-matched non-pregnant rats were inoculated with the following doses of RVFV strain ZH501: 3.1 x 10^3^ pfu (n=11), 3.5 x 10^4^ pfu (n=5), 1.3 x 10^5^ pfu (n=6). Unless the non-pregnant rats met euthanasia criteria and were euthanized, the rats were euthanized at a pre-determined end date of the study at 16 dpi.

Upon necropsy, tissues were harvested and suspended in 2x weight/volume D2 media and homogenized using an Omni tissue homogenizer (Omni International). Tissue homogenates were used to quantitate infectious virus by VPA inside of the BSL-3 facility. For quantitation of RVFV-specific viral RNA by q-RT-PCR analysis, 100 μL of each tissue homogenate was inactivated in 900 μL Tri-Reagent (Invitrogen) for ten minutes prior to removal from the BSL-3 facility. Subsequent storage at −80°C or RNA isolation and q-RT-PCR analyses occurred in a BSL-2 setting.

### RNA isolation and q-RT-PCR

RNA isolation was performed using a modified Invitrogen PureLink Viral RNA/DNA kit protocol. Briefly, 200 μL of chloroform was added to the tissue homogenate/Tri-Reagent, mixed, then centrifuged at 4°C at 12,000 x g for 15 minutes to separate the organic phase from the aqueous phase (contains RNA). The aqueous phase of the cell lysates was collected then mixed with an equal volume of 70% ethanol, then the sample was applied to the PureLink spin column. The PureLink Viral RNA/DNA kit protocol, including DNase treatment, was followed for the remainder of RNA isolation procedure. q-RT-PCR was performed using the SuperScript III Platinum One-Step q-RT-PCR Kit (Invitrogen) following the manufacturer’s guidelines. Primers targeting the RVFV L segment include: RVFV-2912Fwd 5’ TGAAAATTC C TGAGACACATGG 3’ and RVFV-2912Rev 5’ ACTTCCTTGCATCATCTGATG 3’. Taqman probe (RVFV-2950 Probe 5’ CAATGTAA GGGGCCTGTGTGGACTTGTG 3’) were labeled at the 5’-end with the reporter molecule 6-carboxyfluorescein (6-FAM) and quenched internally at a modified “T” residue with BHQ1 (Black Hole Quencher), with a modified 3’-end to prevent probe extension by Taq polymerase (19). Thermocycling parameters include the following: reverse transcription, 50°C for 30 minutes; Taq polymerase inhibitor activation, 95°C for 2 minutes; PCR amplification, 95°C for 15 seconds and 55°C for 30 seconds (40 cycles). Semi-quantitation of virus was determined by comparing CT values from unknown samples to CT values from the in-house developed ZH501 RVFV RNA standards based on PFU equivalents.

### In situ infection of human tissue

Human placental tissue isolated within the second trimester (14-23 weeks) from elective terminations of normal (non-genetically abnormal) pregnancies was obtained from the University of Pittsburgh Health Sciences Tissue Bank through an honest broker system after approval from the University of Pittsburgh Institutional Review Board and in accordance with the University of Pittsburgh anatomical tissue procurement guidelines.

Amnion (fetal membrane), decidua (maternal tissue), and chorionic villi were separated from whole placental tissue and cut to 1 cm x 1 cm sections. Each tissue section was placed in a well of a 24-well plate (Corning) and inoculated with the following doses of RVFV strain ZH501, in duplicate: 1.6 x 107 pfu (n=2 per donor), 3.0 x 10 ^6^ pfu, 3.0 x 10 ^5^ pfu, 3.0 x 10 ^4^ pfu, or 3.0 x 10 ^3^ pfu (n=4 per donor unless otherwise stated). Five hundred microliters of virus diluted in D2 media was added to each tissue and incubated in a 37°C incubator for 1 hour for viral adsorption. The inoculum was removed, washed twice with PBS, then replaced with 1 mL of complete growth media (DMEM/F12, 10% FBS, 1% penicillin-streptomycin, amphotericin B). To generate a viral growth curve, 50 μL was collected from each tissue every twelve hours for forty-eight hours and analyzed by q-RT-PCR. Forty-eight hours post-infection, all supernatant was collected, and tissues were washed two times with PBS then fixed in 4% PFA for 24 hours for fluorescent microscopy imaging. Tissues or supernatant were analyzed in quadruplets; two of these tissues were processed for fluorescent microscopy.

### Fluorescent Microscopy

Human tissue was fixed in 4% paraformaldehyde (PFA) for 24 hours at 4oC, washed in 1x PBS, and then permeabilized with 0.25% Triton X-100 in 1x PBS for 30 minutes at room temperature with gentle agitation. Tissue was washed and then incubated with antibodies to double stranded RNA (recombinant J2, as described previously (20), rabbit anti-cytokeratin-19 (Abcam), and counterstained with actin (using Alexa Fluor conjugated Phalloidin) for 1 hour at room temperature. Following washing with 1x PBS, tissue was incubated with Alexa Fluor conjugated secondary antibodies (Invitrogen), washed, and then mounted with Vectashield (Vector Laboratories) containing 4’,6-diamidino-2-phenylindole (DAPI). Images were captured using a Zeiss LSM 710 inverted laser scanning confocal microscope and contrast adjusted in Photoshop. Quantification of the extent of RVFV infection was performed using Fiji (NIH). Regions of interest (ROIs) of the syncytium from four different donors were defined using Fiji and the extent of RVFV infection (as assessed by the relative fluorescent units; RFU) was quantified. In total, eleven villi were quantified from uninfected controls and sixteen villi were quantified from RVFV infected tissue.

### Histology

For fixation of tissues and inactivation of virus, tissues were submerged for 24 hours in 4% paraformaldehyde (PFA) at 4°C. Prior to removal from the BSL-3 laboratory, the 4% PFA was replaced with fresh 4% PFA. In the BSL-2 laboratory, 4% PFA was removed and washed from tissue with 1x PBS then submerged in PBS and stored at 4°C until processed further. Tissues were paraffin embedded and sections were cut onto slides following standard histological processes by the University of Pittsburgh McGowan Institute Histology Core.

Whole pups were submerged in 4% PFA for 24 hours. To ensure complete inactivation, pups were cut in half laterally, then exposed to 4% PFA for another 24 hours prior to removal from the BSL-3 laboratory. Half pups were further cut into whole head and abdomen sections in the BSL-2 laboratory prior to embedding in paraffin and cut onto slides.

Slides were deparaffinized using an alcohol rehydration series then stained following standard hematoxylin and eosin staining procedures. For immunohistochemistry, fixed slides were deparaffinized using an alcohol rehydration series and boiled in 10mM citric acid buffer pH 6.0 to unmask antigen binding epitopes. Primary antibody was IBT Bioservices Rift Valley Fever MP12 Antibody (Catalog# 04-001); secondary antibody was Vector Laboratories Biotinylated Horse Anti-Rabbit (H+L) (Catalog# BA-1000). Chromogen staining for visualization was carried out using the Vector Laboratories Vector Blue Alkaline Phosphatase Substrate kit according to manufacturer’s instructions. Slides were imaged using the Nikon 90i Eclipse epifluorescent microscope provided by the University of Pittsburgh Center for Biologic Imaging.

### Statistics

Statistical analysis was performed using Graphpad Prism 7.0. Survival of pup and dams was compared using Mann Whitney-U test. Growth curve of pups 7 and 8 from dam 4 was calculated using linear regression modeling. Intensity of RVFV immunostaining in uninfected and infected human placental tissue was compared using a t-test.

## Results

### Susceptibility of late-gestation pregnant Sprague-Dawley rats to RVFV disease and death

Rats develop severe disease after RVFV infection, similar to that observed in domesticated livestock and humans (21, 22). We have previously used rats to study different disease outcomes resulting from RVFV infection (21, 23). Although there are significant morphologic differences between the placentas of humans and rodents, rats are often used to study placental development, embryogenesis, embryotoxicity, and vaccine teratogenicity during pregnancy (24-29). The fully intact placenta forms a physical barrier between the mother and fetus, facilitates nutrient exchange, and protects the developing fetus from microbial invasion. The gestational period for rats is 22 days, and by embryonic day 14 (E14), the placenta is fully formed with deep trophoblast invasion within the uterine wall (24).

For these studies, time-mated Sprague-Dawley (SD) rats at E14 were infected with the pathogenic ZH501 strain of RVFV by subcutaneous injection (s.c.) in the hind flank (Fig. 1A). As controls, age-matched, non-pregnant rats were similarly infected. Some rats developed severe disease as a result of infection, and in these cases, euthanasia endpoint criteria were met within 2-6 days after infection in both pregnant and non-pregnant groups (Fig. 1B). Rats that survived infection had few signs of disease and delivered pups on E22. Surviving pups were not handled until 5 days after delivery, after which they were weighed daily. Surviving dams and pups were euthanized at the end of the study (neonatal age of 10 days; 18 days post-infection (dpi)) (Fig. 1A).

The pregnant rats were more susceptible to death after infection than their non-pregnant counterparts (Fig. 1B). Non-pregnant rats survived infection despite exposure to high doses of RVFV (only 2 out of 22 died; 9%), whereas 57% of pregnant rats died (16 out of 28) when exposed to even low doses of RVFV (p = 0.0008). Within the pregnant rat cohort, there was no association between death after infection and the administered dose (p = 0.321).

### Reproductive tissues are targets of RVFV infection in pregnant and non-pregnant rats

The distribution of virus throughout the tissues was measured by q-RT-PCR and/or viral plaque assay (VPA) in pregnant rats that succumbed to lethal disease (2-6 dpi) (Fig. 1C), pregnant rats that survived (18 dpi; Fig. 1D), and their non-pregnant counterparts (16 dpi for survivors; Suppl. Fig. 1A-B). Infectious virus was widely distributed throughout the animals that succumbed, regardless of pregnancy status. In surviving animals at 18 dpi, infectious viral burden was significantly reduced in both pregnant and non-pregnant rats (data not shown). Viral RNA, however, remained widely distributed in the survivors at the end of the study, 18 dpi (Fig. 1D). As expected because RVFV is a hepatotropic virus, the liver contained large amounts of infectious virus in lethally-infected rats (pregnant and non-pregnant; Fig. 1C & Suppl. Fig. 1B) and high amounts of viral RNA remained in the liver of survivors at 18 dpi and 16 dpi, respectively (Fig 1D & Suppl. Fig. 1A).

Placental samples were available from the pregnant rats that succumbed to disease and a few that survived (most surviving rats consumed the placenta at the time of delivery and were not available for analysis) (Fig. 1E). The placenta contained high levels of infectious virus measured by plaque assay regardless of the time post-infection. Even the placentas obtained from the surviving rats on the day of delivery contained extraordinary amounts of infectious virus, with one animal having over 10^9^ pfu/ml.

In addition to the placenta, the uterus, ovary, and amniotic sac contained infectious virus in the lethally-infected rats (Fig. 1C), and viral RNA persisted in these tissues in survivors (Fig. 1D). Virus was also found in the mammary glands (4/5 rats had detectable vRNA; 1 had infectious virus), indicating vertical transmission of RVFV from dam to pup could possibly occur through lactation. There is indeed a concern that consumption of raw milk from goats and cattle may be a risk for human infection with RVFV (30).

Compared to controls, histology sections of livers from lethally-infected pregnant (Fig. 2) and non-pregnant (Suppl. Fig. 1C) rats displayed a reproducible pattern of tissue injury that included widespread sinusoidal congestion, multifocal recent hemorrhage, massive hepatocyte necrosis, and acute inflammation. These are classic histological findings in animals infected with RVFV (31). In addition, the livers of infected rats had high levels of RVFV-antigen (Suppl. Fig.1 D-E). Altogether, our data suggest that both pregnant and non-pregnant rats succumb to infection following massive liver necrosis, although pregnant rats are more susceptible to developing disease after exposure to low levels of virus compared to non-pregnant rats (Fig. 1B).

**Figure 2:**
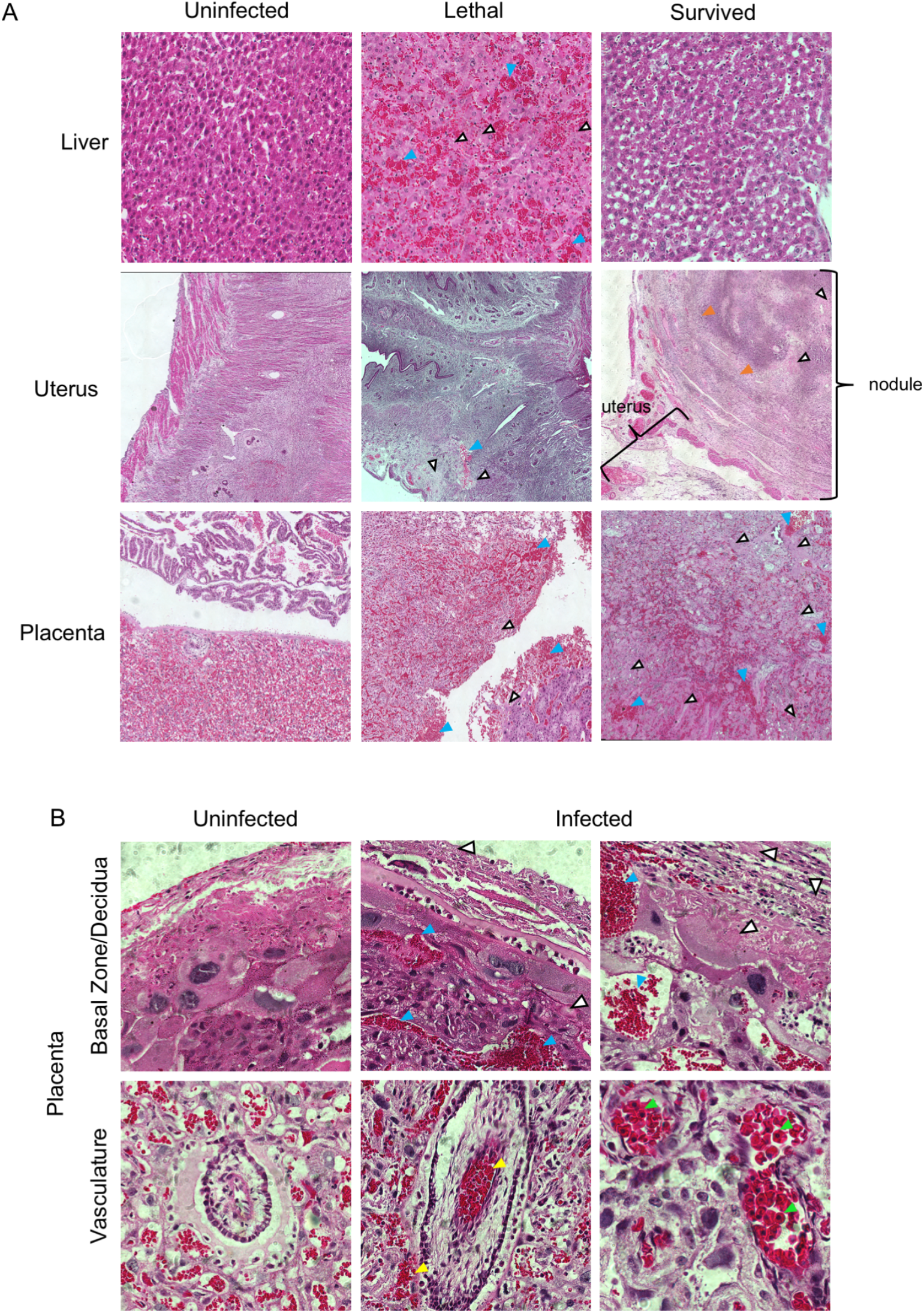
RVFV causes pathology within the liver, uterus and placenta of pregnant dams. H&E staining within the indicated tissues. (A) 20x images of liver, uterus, and placenta. (B) 60x images of placenta. Blue, white, yellow, orange, or green arrow heads highlight evidence of hemorrhaging, necrosis, vascular/perivascular congestion, calcification, or nucleated red blood cells, respectively.

The placentas from lethally and sub-lethally infected dams displayed a reproducible pattern of tissue injury that included multifocal necrosis and recent hemorrhage in the decidua and trophospongiosum; vascular congestion and massive necrosis of parenchymal cells within the labyrinth; and an acute intravascular and perivascular inflammatory infiltrate associated with intraplacental arteries (Fig 2A,B). Also seen was a discernable increase in circulating nucleated red blood cells (nRBCs).

Within lethally-infected dams, necrotic lesions were found within the uterus (Fig. 2A). Several pregnant dams that survived infection had subserosal nodules within the myometrium of the uterus that showed evidence of liquefactive necrosis, acute inflammation, and calcification. Despite the presence of viral RNA and infectious RVFV within the ovaries of both non-pregnant (Suppl. Fig. 1A-B) and pregnant (Fig. 1C-D) RVFV-infected rats, reproducible histopathological changes were not observed.

RVFV vRNA was also found in the testes of infected male Lewis rats (Sup. Fig. 2). The levels of RVFV viral RNA and infectious particles found here in the uterus, ovary, placenta, amniotic sac, mammary glands, and testes, as well as the histological damage in certain tissues, demonstrates that RVFV has a previously unappreciated preference for targeting the reproductive tissues.

### Direct vertical transmission of RVFV to pups during late-gestation results in pup deformity and demise

Within the pregnant dams that succumbed to RVFV infection at 2-6 dpi, virus was widespread throughout various tissues, corroborating the pantropic nature of RVFV (Fig. 1C). Within the corresponding pups from these dams that died, q-RT-PCR of the peritoneal cavity or brains also detected high levels of viral RNA (Fig. 3A). Vertical transmission occurred while the pups were *in utero*, as the dams were euthanized prior to delivery.

**Figure 3:**
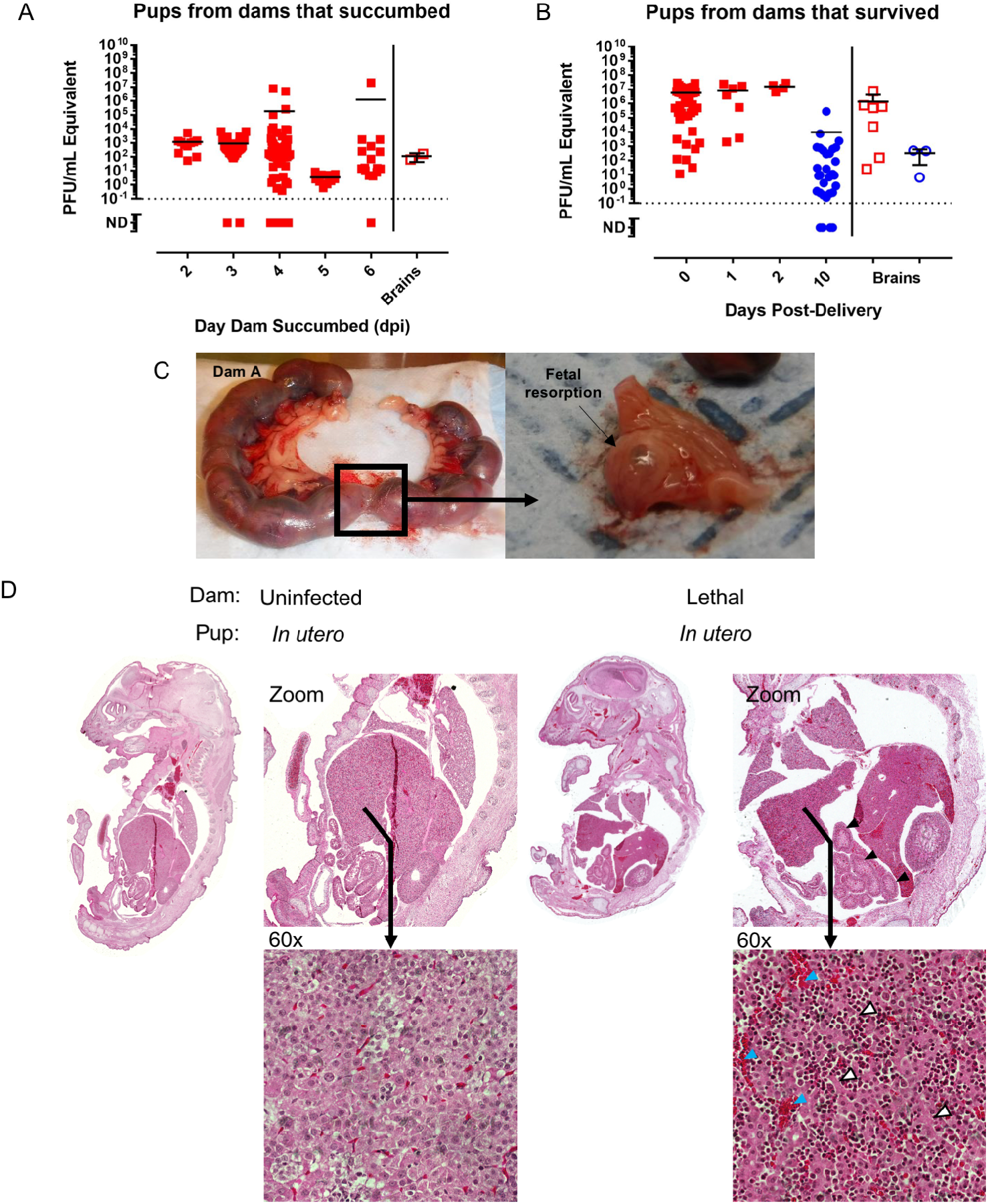
Infection of pregnant dams results in direct transmission of RVFV to the peritoneum and brain of pups. Pups delivered from (A) lethally-infected and (B) surviving pregnant rats were tested for vRNA within the peritoneal cavity (left) and brain (right). In (A), the x-axis represents the day the dams were euthanized due to severe disease. In (B), the x-axis represents the day post-delivery, with day 10 representing surviving pups euthanized at the end of the study. For both graphs, red square data points indicate pup demise and blue circle data points indicate pup survival. Open data points are pup brain tissues; all closed data points are pup peritoneal cavity. (C) Photographic evidence of fetal resorption within the uterus of one of three dams that succumbed to RVFV infection. (D) 10x images of whole pups were examined for histological changes. H&E staining of a whole pup from a dam that succumbed to infection (right) or a corresponding uninfected control rat (left) euthanized at the same day of gestation. Blue, white, and black arrow heads highlight evidence of hemorrhaging, necrosis, or altered intestinal structure, respectively.

Because most humans develop non-lethal febrile illness when infected with RVFV, we were interested in the outcome of the pups from dams that survived infection with few to no clinical signs. Remarkably, many of the pups born from these surviving, clinically-normal dams were dead at delivery or shortly thereafter (Fig. 3B). Of ten pregnant dams that survived infection and reached full-term (E22), all gave birth to at least one pup that died either due to infection or consumption on the days following delivery. An additional dam survived to full-term; however, all pups were consumed by this dam and were unable to be recorded for analysis.

The eleven infected dams gave birth to a total of 111 pups, 39 of which survived (35.1%). In comparison, of the eight uninfected control dams, 66 of 88 (75.0%) pups survived to the end of the study (p < 0.0001). Dams are known to eat their young due to stress (such as the laboratorian disturbing the pups to weigh them after delivery) or if they notice abnormalities (17, 18). Some of the uninfected dams consumed their pups during this study, so the survival rate of the pups from uninfected dams can be considered the baseline for these experimental conditions. The average survival rate of pups per litter delivered by infected dams was 27.2%, whereas uninfected rats were 2.45-times more likely to survive (66.8%; p = 0.02). Outcomes from 4 specific dam cases are presented and discussed in the following section.

Overall, we detected very high levels of viral RNA (106-107 pfu/mL eq.) in the peritoneal cavity of still-born pups birthed by dams that had no apparent clinical disease (Fig. 3B). It is quite striking that the virus was vertically transmitted and replicated to high levels in the pups, while causing no clinical disease in the dam herself. Viral RNA was also detected in the brains of pups from both surviving and lethally-infected dams (Fig. 3A,B). Fetal resorption is characteristic of vertical transmission in animals infected with viruses such as Zika virus (32) and vesicular stomatitis virus (33). Evidence of fetal resorption occurred in three lethally-infected dams exposed to RVFV during late-stage gestation (Fig. 3C & Suppl. Fig. 3).

H&E sections of whole pups from lethally-infected dams showed significant liver damage (massive parenchymal necrosis, vascular congestion and areas of recent hemorrhage) (Fig. 3D). Pups also displayed altered intestinal structure compared to pups from uninfected dams.

During the late stages of pregnancy, RVFV is directly transmitted to the developing pups resulting in increased rates of death and tissue damage. Interestingly, this is not an all-or-nothing phenomenon within individual litters, as we found that physiological changes and survival varied within the same litter. To illustrate this, four selected cases are described below.

### Description of individual dam cases

Dam 1 reached full-term gestation with no apparent clinical signs nor significant liver damage, yet she still died giving birth (8 dpi). Dam 1 delivered eleven pups that were all dead upon delivery, with 2 pups remaining within the uterus at the time of her demise (Fig. 4A). Like other RVFV-infected dams, viral RNA was widespread throughout the dam’s tissues (Fig. 4B). Because this dam died during delivery, the placenta was available to directly compare the virus burden of the placenta to other maternal tissues harvested at the same time. In the other cases discussed below, the dams ate placentas after birth, so they were unavailable to sample. Strikingly, at the time of death, dam 1 had 107 pfu/mL of virus in the placenta, which is 4-logs more viral RNA than the liver and spleen, the primary targets of the virus (Fig. 4B). In addition, the uterus and ovary had about 1-log more viral RNA than the liver and spleen, demonstrating a preference for RVFV to replicate within reproductive tissues.

**Figure 4:**
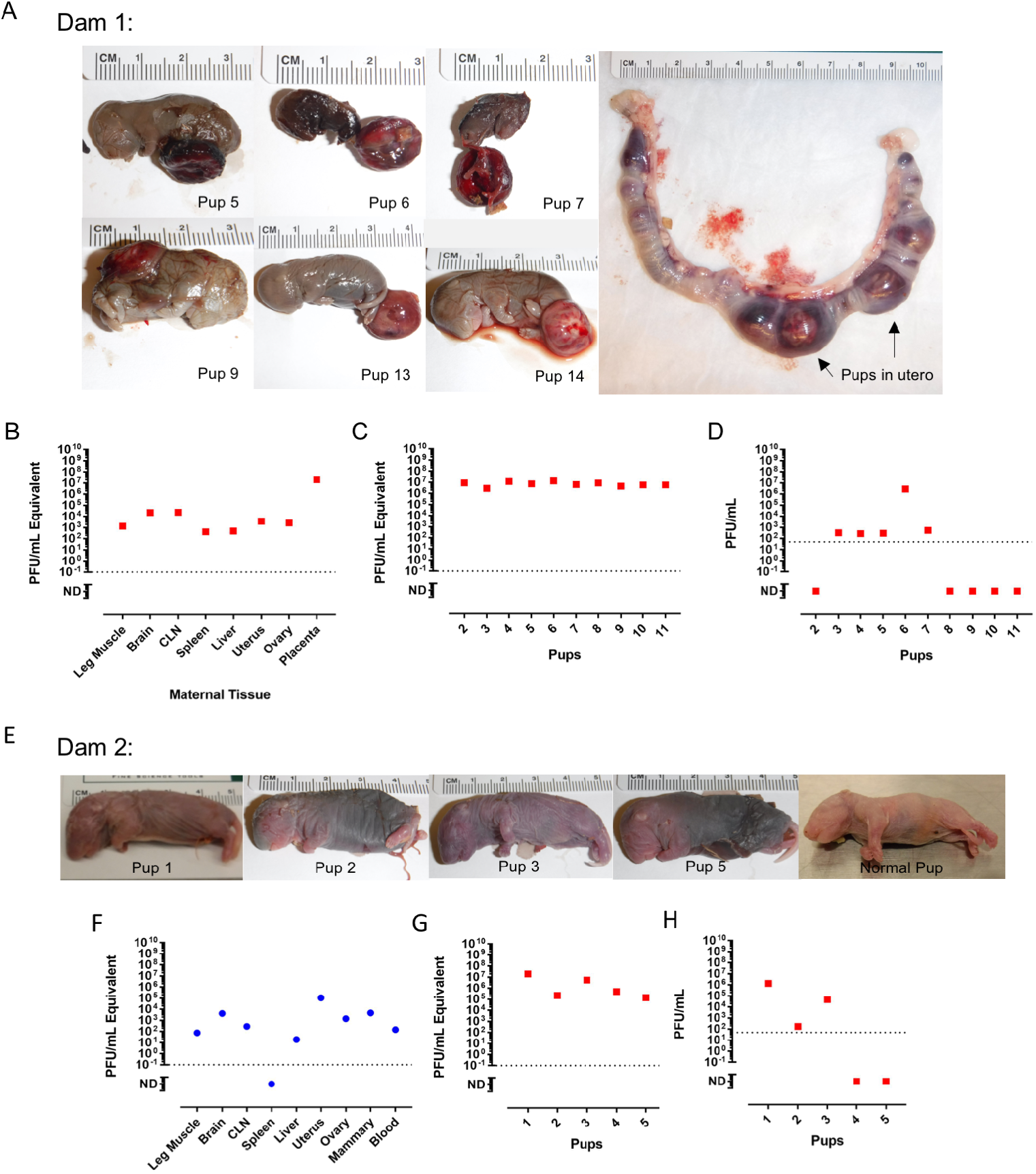
Vertical transmission of RVFV in 2 dams resulted in still-born pups with physiological abnormalities and high viral titers. (A-D) Dam 1 was inoculated with 175 PFU and died giving birth at 8 dpi. Dam 1 delivered 13 still-born pups, with 2 remaining in the uterus. (A) Pictures of individual pups are shown as numbered. Uterus with pups 13 and 14 within birth canal post-demise. (B) vRNA within maternal tissue. (C) vRNA and (D) infectious virus within peritoneal cavity of indicated pups. The three pups not included in the graphs were used for other analyses. (E-H) Dam 2 was inoculated with 175 PFU and survived with no clinical signs of disease. Dam 2 delivered 7 pups to full-term; 5 were dead and 2 were alive but subsequently consumed by the dam (data not available). (E) Pictures of individual pups as labeled. A normal pup from an uninfected dam is shown for comparison. (F) vRNA within maternal tissue. (G) vRNA and (H) infectious virus within peritoneal cavity of indicated pups. For all graphs, red square data points indicate either dam or pup demise. Blue circle data points indicate dam or pup survival.

The stillborn pups from dam 1 had apparent gross abnormalities (small, rounded head and grey discoloration) and variable size (1.5 – 3 cm) compared to uninfected pups (5 cm; Fig. 4A). Nine of the eleven pups (82%) had abnormally shaped heads compared to uninfected pups. Pup 6 and 7 likely died earlier *in utero*, as indicated by their small size and apparent decomposition; both of their heads were disproportionately large due to early gestational demise. While the other pups were larger than pups 6 and 7, none of them were normal in size, and they were all born deceased. The pups had very high levels of viral RNA (10^6^—10^7^ pfu/mL eq., irrespective of pup size; Fig. 4C). Less infectious virus was recovered from the pups compared to viral RNA (Fig. 4C,D), likely due to the inability to preserve tissue immediately after death because pups succumb to infection *in utero* or shortly after delivery. This trend is observed for all representative dams (Figs. 4-5), otherwiseRVFV infectivity correlates very closely with q-RT-PCR results as seen in Fig. 1.

Dam 2 survived infection with no apparent clinical signs. Viral RNA was still detected throughout the dam at 18 dpi, with the highest burden in reproductive tissues and the brain (Fig. 4F). At E22, she delivered seven pups; five were born dead and two were alive. Both living pups were consumed by the dam within a day of birth. Dams periodically eat their young due to stress or if they notice abnormalities in their pups (17, 18). The pups shown in Fig. 4E are around 5 cm in length, which is normal for full-gestation pups. Several have dark colored bodies likely signifying decomposing tissue and/or blood congealment. Smaller, rounded heads without a pointed snout were observed in three of the seven pups (pups 2, 3, and 5). Despite pup 1 having similar physiological features and length as a normal pup delivered from an uninfected mother (Fig. 4E), viral RNA and infectious virus was highest in this pup compared to its littermates that had more severe gross anatomical abnormalities (Figs. 4G-H). This may be due to more advanced decomposition setting in pups 2, 3, and 5 that affected virus infectivity.

In another case, dam 3 also survived infection with no clinical signs of disease and had widespread viral RNA in tissues at 18 dpi (Fig. 5B). She delivered ten pups at full-term (E22); six pups were born deceased and four were alive. Two of the living pups survived until the scheduled euthanasia date (18 dpi; 10 days neonate), whereas the other two were consumed by the dam within two days post-delivery. Pups 1 and 2 from dam 3 were relatively normal in appearance; however they were born deceased and displayed the highest viral RNA (Fig. 5C) and infectious virus (Fig. 5D) burden compared to littermates with physiological abnormalities; this could be due to the more recent demise of pups 1 and 2 which preserved infectious virus until tissue was collected. The pups who die earlier *in utero* likely have less infectious virus due to decomposition which affects virus viability. Despite pups 7 and 8 surviving until planned euthanasia at neonatal age of 10 days with no signs of disease or gross abnormalities, viral RNA was still detected in the peritoneal cavity of these pups, although infectious virus was not found (Figs. 5C-D). Pup 7 weighed significantly less than its littermate, and both had similar growth rates (slope = 1.88 for pup 7 and 2.26 for pup 8; p = 0.14) after birth Fig. 5E). It is unclear if this is natural variation in pup size or an effect of viral infection *in utero*.

**Figure 5:**
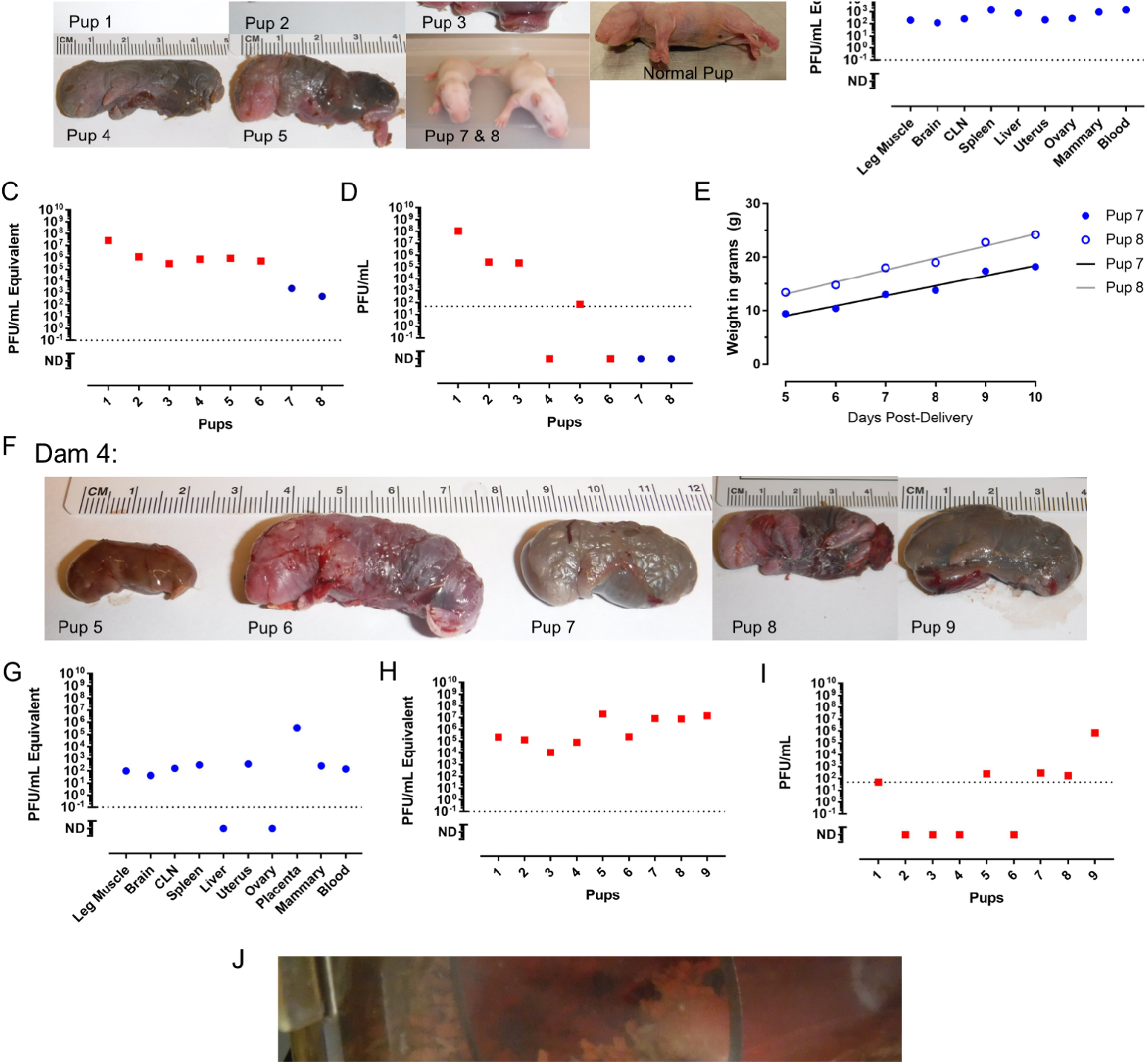
Evidence of variable survival and physiological outcome of pups resulting from vertical transmission. (A-E) Dam 3 was inoculated with 175 PFU and survived with no clinical signs of disease. Dam 3 delivered 10 pups to full-term; 6 dead and 4 alive. Two living pups were consumed by the dam within 2 days of birth (data not available). Two remaining pups survived to the end of the study (pups 7 and 8). (B) vRNA within maternal tissue. (C) vRNA and (D) infectious virus within peritoneal cavity of indicated pups. (E) Weight of surviving pups from dam 3 from days 5 to 10 neonate. Black or grey lines represent growth curve calculated by linear regression modeling of pup 7 or 8, respectively. (F-J) Dam 4 was inoculated with 1300 PFU and survived with no clinical signs of disease. Dam 4 delivered 9 still-born pups. (F) Pictures of individual pups as labeled. (G) vRNA within maternal tissue. (H) vRNA and (I) infectious virus within peritoneal cavity of indicated pups. (J) Cage of dam 4 as it was found on day of pup delivery (E22; 8 dpi).

Finally, dam 4 had no signs of clinical disease and delivered a total of nine deceased pups at full-term (Fig. 5F). Multiple pups were found encapsulated in an amniotic sac and/or still connected to a placenta (Fig. 5J). Placenta collected the day of delivery had high levels of viral RNA (10^5^ pfu/ml eq.); the viral RNA levels in the placenta were similar to that found in the pups, regardless of pup size (Figs. 4H,I). Seven out of nine pups (78%) had abnormally shaped heads compared to uninfected pups. The pups had gross abnormalities and varied in color from pink to gray to black. Infectious virus was detected in five out of nine pups (Fig. 4I). The smallest pup (pup 11), measured at about 2.2 cm, while the largest pup from the litter was about 4.1 cm in length; all pups from this litter were still smaller than pups delivered from uninfected dams (typically 5 cm).

The four dams highlighted above, despite being asymptomatic, all had liquefactive myonecrosis within the uterus (Fig. 2). The placenta also had striking abnormalities, including inflammation, cell necrosis, and hemorrhage in the decidua and villous structures (Fig 2A-B).

### Vertical transmission of RVFV in an early-gestation pregnant rat

To examine the potential for vertical transmission of RVFV prior to the development of a fully formed placenta, an early-gestation (E5) Sprague Dawley dam was infected with 1.5 x 10^5^ pfu of RVFV along with an uninfected, age-matched, pregnant dam as a control. At seven days post-infection, corresponding to E13, neither dam had clinical signs of disease and both dams underwent planned euthanasia (Suppl. Fig. 4A). The observed virus distribution in the early-gestation pregnant rat resembled that of the late-gestation pregnant rats (Suppl Fig. 4B and Fig. 1). Infectious virus was detected in the cervical lymph node, spleen, liver and placenta, with the placenta containing 3-logs more infectious virus than the spleen (10^6^ vs 10^3^ pfu/mL, respectively; Suppl. Fig. 4B). Due to the small size of each embryo at this gestational age, embryonic tissue from the entire litter was pooled and analyzed for the presence of viral RNA and infectious particles (Suppl. Fig. 4C). Pooled embryos had 2-log more viral RNA and infectious virus than the spleen of the dam (10^6.5^ vs 10^3^ pfu/mL respectively).

At the time of euthanasia, the uninfected dam had a healthy uterus containing fourteen embryos, all with similar sizes, shapes, and a translucent, pale-yellow coloring. Of the sixteen embryos of the infected dam, four (25%) were red in color (Suppl. Fig. 4D) instead of the pale-yellow coloring observed in the remaining twelve embryos and embryos of the uninfected control (Suppl. Fig. 4D, left). Three embryos were smaller than the embryos from the rest of the litter. They lacked amniotic fluid and were palpably dense and hard in appearance (Suppl Fig. 4D; embryos 4-6); embryos 4 and 5 were two of the four red in color. The red coloring and rigid appearance and structure of embryos from infected dams suggests early hemorrhaging and/or resorption of the embryos. These results show that RVFV is vertically transmitted to embryos of dams infected during early-gestation, with a preferential niche within the placental tissue.

### Replication of RVFV in human placental tissue

Although two isolated cases of vertical transmission have been reported in humans from Africa and the Middle East (13, 14) and a single study highlighted the potential of RVFV-infected women to be 4-times more likely to have a late miscarriage or stillbirth (15), the effect of RVFV infection on the developing human fetus is not known. To evaluate whether human fetal tissue is permissive to RVFV infection and replication, placental chorionic villi were obtained from healthy human donors undergoing elective termination at 16-23 weeks of gestation (second trimester). Tissues were inoculated *in vitro* with RVFV, and explant supernatant was collected every 12 hours for measurement of viral replication. Similar viral growth kinetics was seen in chorionic villi from 2 different donors (Fig. 6A); 2-3 log increase in virus production was detected between 12 and 36 hours post-infection.

**Figure 6:**
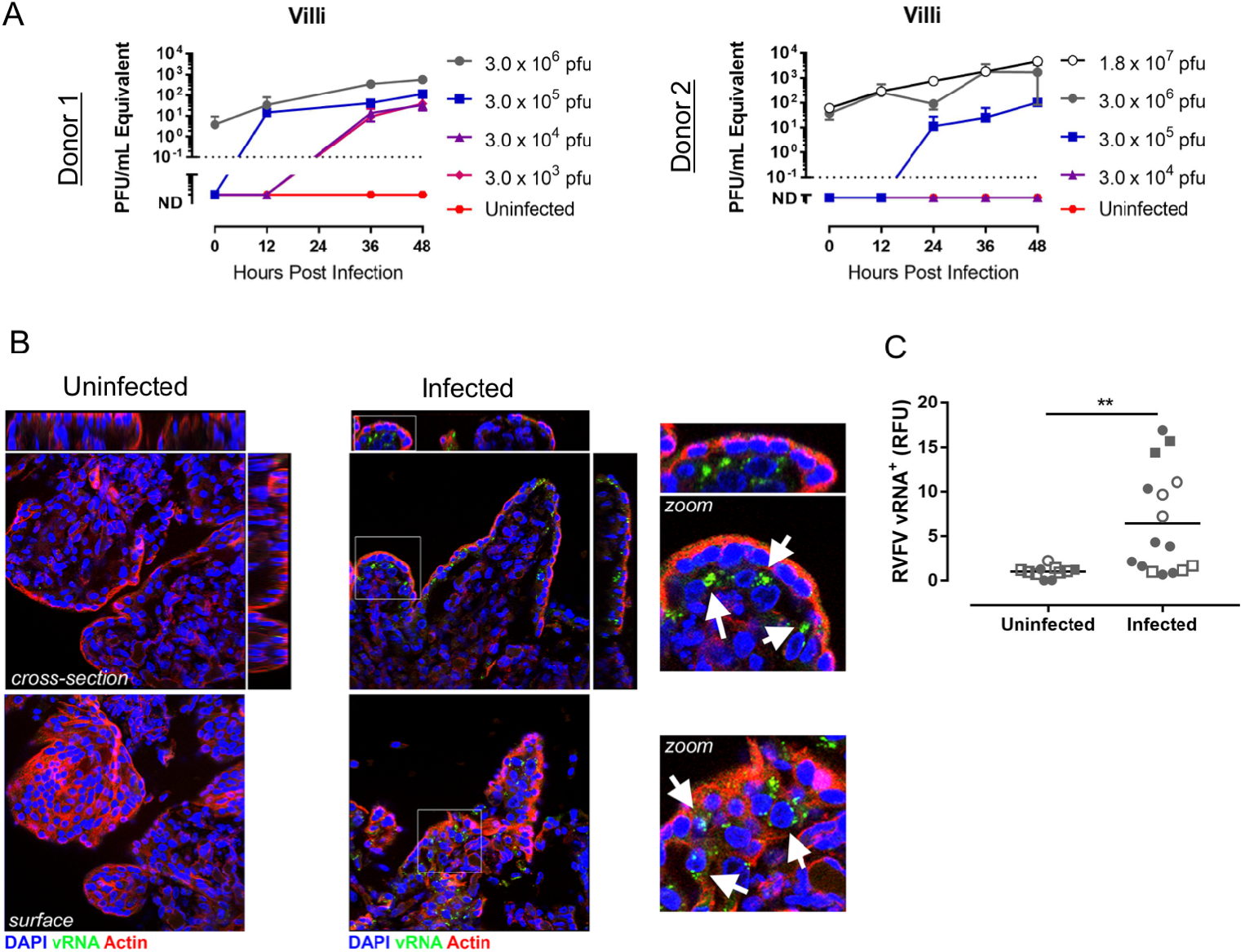
RVFV replicates in human placental tissue, even the highly resistant syncytiotrophoblasts within placenta villi. Human chorionic villous tissue explants from donors 1 and 2 (A) were infected in vitro with RVFV at the indicated doses, then supernatant was harvested at 0, 12, 24, 36 and 48 hours post-infection for measurement of vRNA by q-RT-PCR. (B) Immunofluorescent microscopy images of villi infected with 3 x 10^6^ pfu RVFV for 48 hours. Uninfected control shown in left panels. DAPI (blue) stains DNA, J2 antibody (green) stains dsRNA of RVFV, cytokeratin 19 (red) stains epithelial cells, actin (purple) stains all cells. (C) Fluorescent dsRNA signal was quantified from 4 human donors. Data from uninfected (n=11) and infected (n=16) villi are shown with symbols stratified by donor (close circles, donor 1; open circles, donor 2; closed squares, donor 3; open squares, donor 4).

To identify the cell type(s) targeted by RVFV in placental villi, we performed immunostaining for double-stranded RNA as described (20). Viral replication, as assessed by the production of dsRNA, was evident in the villi, with replication observed in both the syncytial layer and the subsyncitial layers within mononuclear cytotrophoblasts (Fig. 6B). Significantly higher levels of dsRNA signal was detected in infected villi from four different donors compared to uninfected villi from the same donors (Fig. 6C). These data suggest that RVFV exhibits tropism for human placental tissue and can replicate to some degree in syncytiotrophoblasts which are known to resist many viruses (34).

## Discussion

Viral infections in pregnant women can have adverse effects on both the mother and the developing fetus. Pregnant women are likely to have more severe pathology, more serious complications, and an increased risk of death compared to non-pregnant women (35, 36). For both seasonal and pandemic influenza, pregnant women are more susceptible to developing severe complications requiring hospitalization (37, 38). Other emerging viruses, like Ebola and Lassa, are also more severe or deadly in pregnant women (39, 40). Here, we found pregnant rats to be more susceptible to severe disease and death after RVFV infection than their non-pregnant counterparts. Necrosis of the liver was characteristic of disease in both pregnant and non-pregnant rats, indicating that pregnancy did not alter the disease presentation, but rather made the pregnant animals more susceptible to severe disease overall.

A 1987 study of pregnant women in Mozambique showed that women with miscarriage or stillbirth had the same prevalence of RVFV-specific IgG antibodies as women with normal deliveries (41). However, detection of IgG antibodies is indicative of past exposure to RVFV and not necessarily acute infection during pregnancy. In a more recent cross-sectional study in Sudan, there was a significant association between acute RVFV infection during pregnancy and miscarriage (15). Of the women with confirmed RVFV infections and disease during pregnancy, 54% had miscarriage compared to 12% in uninfected women. Acute RVFV infection was an independent predictor of miscarriage (odds ratio of 7.4); all of these miscarriages occurred in the 2^nd^ and 3_rd_ trimesters (15). While this is only one study, it strongly implicates RVFV infection as a causative agent of miscarriage in late-gestation. Miscarriage earlier in pregnancy due to RVF may also occur but remain underreported.

Two cases of vertical transmission of RVFV were documented in Saudi Arabia (2000) and Sudan (2007) (13, 14). In both cases, onset of RVF-like illness in the mothers occurred around 37-38 weeks of gestation. In the two weeks prior to illness in the Saudi woman, 6 family members developed RVF-like illness and 1 died of confirmed RVF. The woman developed RVF-like symptoms 4 days prior to delivery of a full-term infant at home. On the second day of life, the infant was hospitalized with respiratory distress, jaundice, and lethargy. He died 6 days after admission with hemorrhagic signs and an enlarged liver. In the Sudanese patient, labor commenced within 10 days of illness onset, and an infant was born with skin rash, palpable liver and spleen, and an Apgar score of 5. Samples from the mother and cord blood tested positive for RVFV IgM. The outcome of this infant is not known. Taken together, these data suggest that acute RVFV infection is associated with adverse pregnancy outcomes, specifically vertical transmission during the second and third trimesters.

To our knowledge, this is the first study to discover the degree to which RVFV targets reproductive tissues and causes catastrophic pathology. The ovaries, uterus, and placenta are previously unrecognized sites of virus replication in both pregnant and non-pregnant immunocompetent animals. A previous study in mice found virus-infected macrophages within the stroma of ovaries from immunodeficient animals infected with an attenuated strain of RVFV (42). Here, we not only found high levels of infectious virus and viral RNA in the reproductive tissues, but the placenta and uterus displayed histologic changes associated with viral infection. Liquefactive necrosis within the uterus of the four dams profiled was striking. Uterine cell death may be a contributing factor to premature placental detachment and intrauterine fetal death within RVFV infected dams. Additionally, the tissues with the highest viral burden were the liver and placenta of lethally and sub lethally-infected pregnant dams. Similarities in physiological functions of these two organs and the high viral load and pathology (massive vascular congestion, recent hemorrhage, cell necrosis, and acute inflammation) observed during infection suggest similar mechanism of disease and infection. Further studies should be performed to understand the onset and progression to fetal demise and elucidate commonalities between the liver and placental structure that accommodates comparable pathology after RVFV infection.

The most alarming and arresting finding from our study was the delivery of dead pups from dams that survived RVFV infection, appeared otherwise clinically normal, and gave birth at normal gestation length. The pups displayed physical abnormalities that resembled *hydrops fetalis*, which is an abnormal accumulation of fluid in the fetus. *Hydrops fetalis* is seen in RVFV-infected livestock (43) and has been noted as a prominent occurrence in pregnant women infected with parvovirus and Zika (44-47). Most humans infected with RVFV develop mild febrile disease, and our study suggests that mild infection of pregnant women may still have devastating impacts on the developing fetus.

The data presented also demonstrates the heterogeneous survival and physiological outcomes of pups delivered by RVFV-infected dams. Some dams gave birth to both live and dead pups, and within the dams that delivered only dead pups, there was variation in the size and gross abnormalities seen within pups from the same litter, indicating that infection, growth restriction, developmental abnormalities, and/or death did not occur simultaneously or with the same outcome. Further studies will determine the specific developmental effects that RVFV infection *in utero* has on the developing fetus.

Our work in pregnant rats pointed to the severe teratogenic effects of RVFV. However, given that the human and rodent placenta differ at both the morphologic and cellular levels, which may influence vertical transmission, we also performed studies in human tissue isolated from mid-gestation. Importantly, this gestational age represents a stage in which RVF-induce fetal death has been observed in humans (15). Remarkably, our studies revealed that RVFV can replicate to a high degree in human chorionic villi, which resist infection by other viruses, including ZIKV (20,48,49). Immunofluorescence analysis showed that viral replication occurred in both the outermost syncytial layer as well as in subsyncytial layers such as cytotrophoblasts. Of note, we have shown previously that human placental syncytiotrophoblasts are highly resistant to viral infections through their constitutive release of antiviral molecules including type III interferons (IFNs) (48,50,51). We have also shown that the syncytium expresses high levels of interferon stimulated genes (ISGs) under basal states, suggesting that these IFNs protect the syncytium from viral infections in an autocrine manner. Our data thus suggest that RVFV is insensitive to placental-derived type III IFNs, perhaps through mechanisms that suppress the activity of ISGs or the response of the syncytium to type III IFNs. Future studies aimed at defining these mechanisms will be critical to design therapeutic approaches to reduce the RVFV vertical transmission.

Ruminants have multicotyledonary, epitheliochorial placentas while humans and rodents both have discoid hemochorial placentas (24, 52). Despite some similarities, the placental structure differs between rats and humans in many ways. Humans have a placental-fetal interface consisting of villous trophoblasts with extravillous trophoblastic cells, while rats have a labyrinth zone with invasive trophoblasts (24). Although livestock have different placental structures than rodents and humans, our data suggests that the mechanism by which RVFV is vertically transmitted can be shared amongst eutherian mammals, with specific tropism for fetal-derived placental cells. Our study thus provides a framework for understanding vertical transmission of RVFV that is applicable to both a veterinary and human context. Comparison of these models could also explain the potentially disparate rate of miscarriages between livestock and humans.

While the abortogenic effect of RVFV in pregnant livestock is historically well-known, the mechanisms underlying these observations are not. The teratogenic potential of live-attenuated vaccine candidates are assessed in pregnant sheep, despite the lack of data on the basic mechanism of vertical transmission of wild-type virus (53, 54). Our findings here are noteworthy because this is the first time direct vertical transmission through the placenta has been demonstrated in a species other than livestock. Because the ability to test vaccines in livestock is expensive and limited to only a few facilities able to perform these types of experiments, evaluation of teratogenicity in rats would be very useful. Vaccine candidates could be screened for adverse effects in pregnant rats prior to evaluation in pregnant ewes.

This study provides insights into the teratogenic potential of RVF in humans and highlights the need for more epidemiological data from human outbreaks to understand the effect of acute RVFV infection in pregnant women. The data that exist, combined with what is known to occur in livestock and our work presented here, all point to the high likelihood that RVFV can have damaging effects on fetuses in humans. Emergence of RVFV beyond its current locations due to changing climate, altered mosquito habitats, and/or accidental introduction would present a significant risk to pregnant women. Future work aimed at defining teratogenic potential of RVF in humans is critical in order to design strategies to reduce the potential for fetal disease in pregnant women, who may themselves display few clinical symptoms.

## Acknowledgements

The authors would like to express sincere gratitude for the technical assistance provided by Aaron Walters, advice on experimental design from Jeneveve Lundy and Reagan Walker, rat art provided by Henry Ma, and study coordination by Stacey Barrick. We also thank the Center for Biologic Imaging and the McGowan Center for Regenerative Medicine for histology support. This work was partially supported by the University of Pittsburgh Center for Vaccine Research (A.L.H), NIH R01-NS101100-01A1 (A.L.H), R21-NS088326 (A.L.H), NIH R01-AI081759 (C.B.C.), R01-HD075665 (C.B.C.), a Burroughs Wellcome Investigators in the Pathogenesis of Infectious Disease Award (C.B.C), and the Children’s Hospital of Pittsburgh of the UPMC Health System (C.B.C.). The authors would also like to acknowledge the Tissue and Research Pathology Services/Health Sciences Tissue Bank, which receives funding from P30CA047904.

## Author Contributions

C.M.M, C.B.C, and A.L.H conceived the study and designed the experiments; C.M.M., N.A., D.A.B., J.R.A, and M.R.K. performed experimental work; J.F.B contributed pathology expertise and interpretation; C.M.M., C.B.C, and A.L.H. analyzed and interpreted the data; C.M.M. and A.L.H wrote the manuscript.

---

**Supplemental Figure 1:**
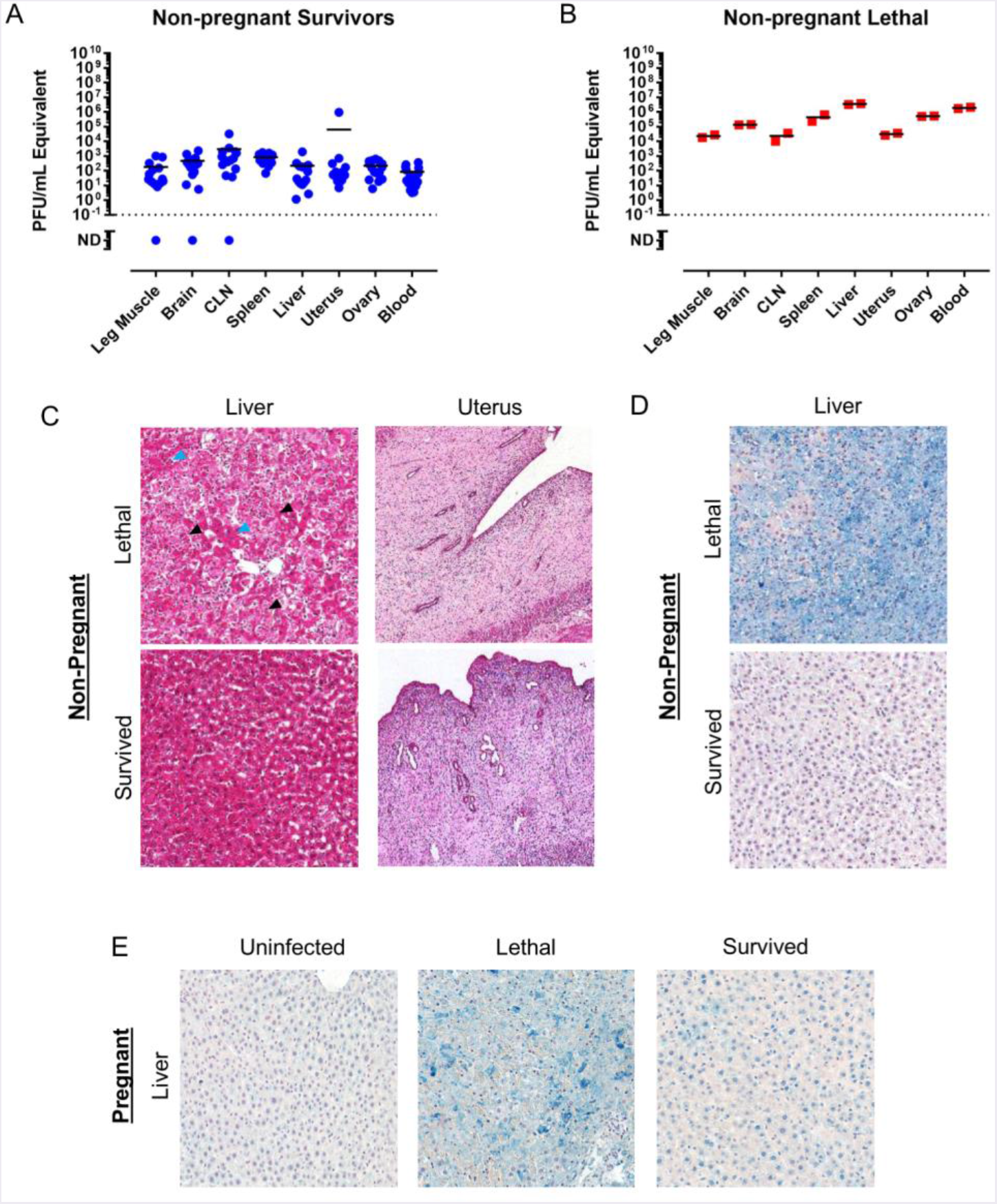
RVFV is present in the ovary and uterus of non-pregnant female rats. Tissues from non-pregnant female rats shown in Fig. 1B were tested for vRNA by q-RT-PCR. Rats that (A) survived (n=15) or (B) succumbed (n=2) to infection. (C) H&E staining or (D,E) chromogen staining (blue) for RVFV antigen in the indicated tissues. (C,D) tissues from infected non-pregnant rats that survived (18 dpi) or succumbed to infection (2 dpi). (E) RVFV antigen staining of liver from uninfected, lethal (4 dpi), or surviving pregnant dams (18 dpi; E14). Black or blue arrow heads highlight evidence of necrosis or hemorrhaging. 20x objective images.

**Supplemental Figure 2:**
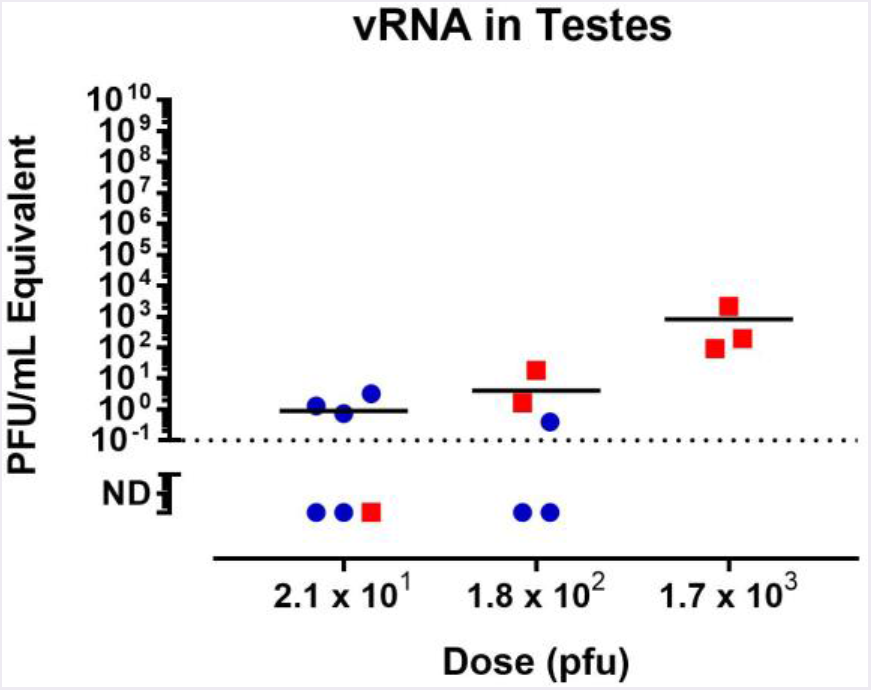
Detection of RVFV in the testes of male rats. Male Lewis rats were infected with RVFV (s.c.). Testes samples were taken from rats that succumbed to disease (red squares) and those that survived (blue circles) and tested for vRNA by q-RT-PCR.

**Supplemental Figure 3:**
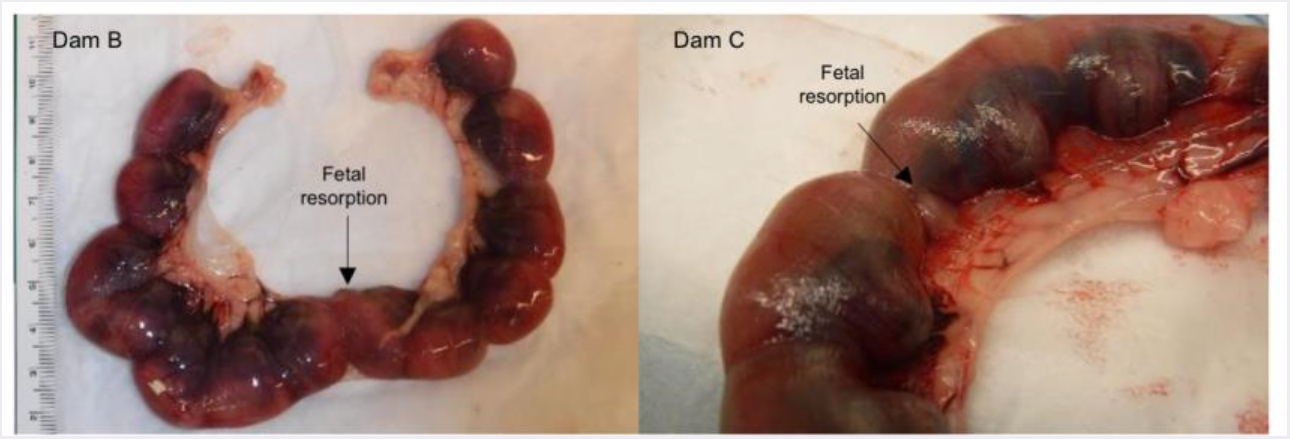
RVFV infection leads to fetal resorption. Photographic evidence of fetal resorption within the uterus of two of three dams who succumbed to RVFV infection.

**Supplemental Figure 4:**
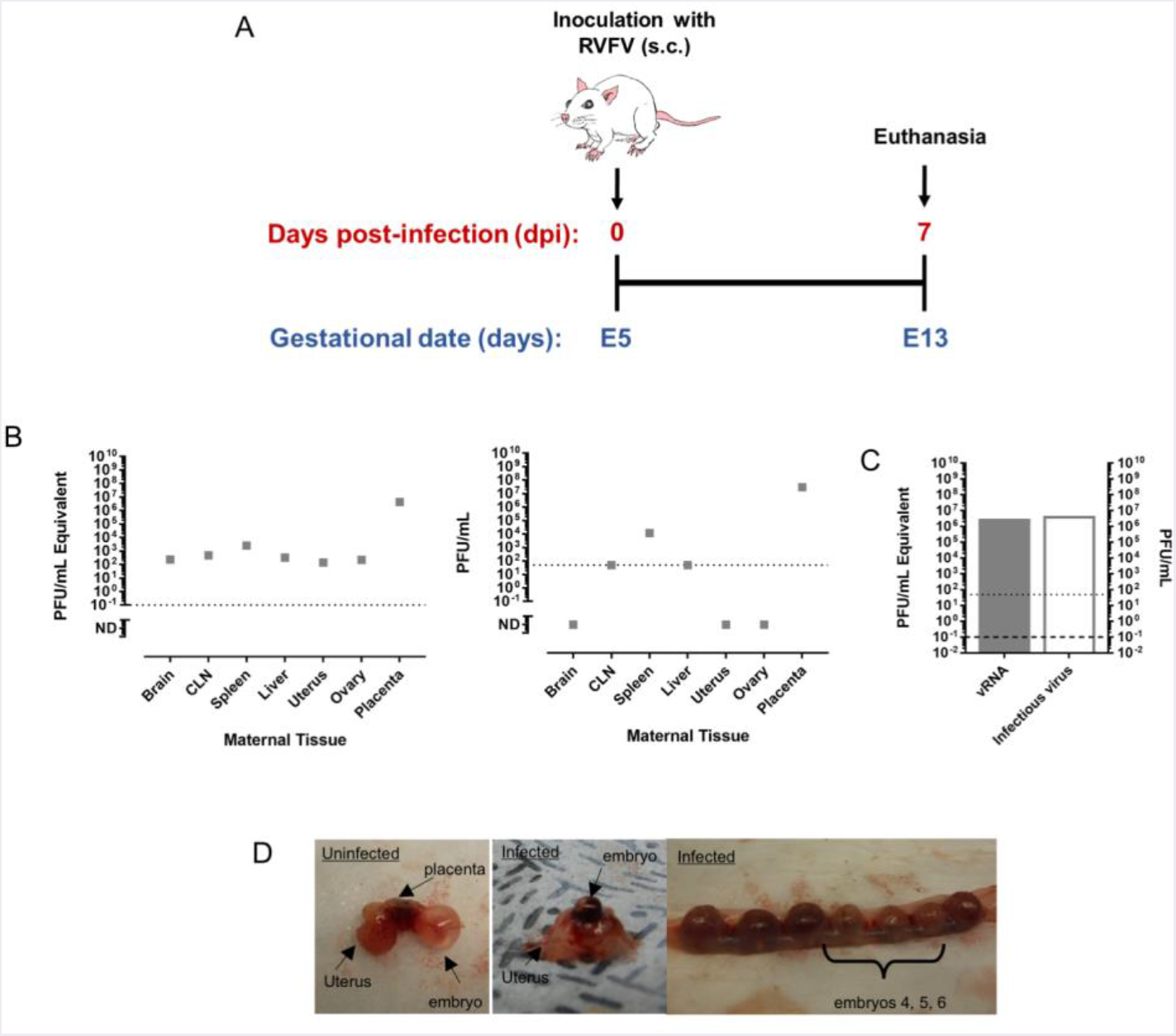
Evidence of vertical transmission of RVFV in an early gestation (E5) pregnant dam, with high viral titers in the placenta. (A) An early gestation (E5) dam was infected with 1.46 x 10^5^ pfu (s.c.) and then euthanized at the pre-determined date of 7 dpi (E13) with no signs of disease. (B) vRNA (left) and infectious virus (right) within maternal tissue. (C) vRNA (left y-axis, solid grey bar) and infectious virus (right y-axis, open grey bar) within pooled embryos (16 embryos). (D) Photographic evidence of embryos from uninfected or infected dams at E13. Uterus containing embryos from an infected dam (right panel).

## References

1. LaBeaud AD, Pfeil S, Muiruri S, Dahir S, Sutherland LJ, Traylor Z, Gildengorin G, Muchiri EM, Morrill J, Peters CJ, Hise AG, Kazura JW, King CH. 2. Factors associated with severe human Rift Valley fever in Sangailu, Garissa County, Kenya. PLoS neglected tropical diseases. 9(3):e0003548. PMCID: PMC4357470.

2. Anyangu AS, Gould LH, Sharif SK, Nguku PM, Omolo JO, Mutonga D, Rao CY, Lederman ER, Schnabel D, Paweska JT, Katz M, Hightower A, Njenga MK, Feikin DR, Breiman RF. 2010. Risk factors for severe Rift Valley fever infection in Kenya, 2007. Am J Trop Med Hyg. 83(2 Suppl):14–21. PMCID: PMC2913492.

3. Smithburn KC, Haddow AJ, Lumsden WH. 1949. Rift Valley fever; transmission of the virus by mosquitoes. Br J Exp Pathol. 30(1):35-47. PMCID: PMC2073112.

4. Linthicum KJ, Britch SC, Anyamba A. 2016. Rift Valley Fever: An Emerging Mosquito-Borne Disease. Annu Rev Entomol. 61:395–415.

5. Turell MJ, Britch SC, Aldridge RL, Kline DL, Boohene C, Linthicum KJ. 2013. Potential for mosquitoes (Diptera: Culicidae) from Florida to transmit Rift valley fever virus. Journal of Medical Entomology. 50(5):1111–7. PMCID: PMCPMID: 24180117.

6. Brustolin M, Talavera S, Nunez A, Santamaria C, Rivas R, Pujol N, Valle M, Verdun M, Brun A, Pages N, Busquets N. 2017. Rift Valley fever virus and European mosquitoes: vector competence of Culex pipiens and Stegomyia albopicta (= Aedes albopictus). Med Vet Entomol. 31(4):365–72.

7. Vloet RPM, Vogels CBF, Koenraadt CJM, Pijlman GP, Eiden M, Gonzales JL, van Keulen LJM, Wichgers Schreur PJ, Kortekaas J. 2017. Transmission of Rift Valley fever virus from European-breed lambs to Culex pipiens mosquitoes. PLoS neglected tropical diseases. 11(12):e0006145. PMCID: PMCPMC5760105.

8. World Health Organization. Annual review of diseases prioritized under the Research and Development Blueprint. 2017 January 24-25, 2017. http://www.who.int/blueprint/what/research-development/2017-Prioritization-Long-Report.pdf?ua=1.

9. Easterday BC. 1965. Rift Valley Fever. Advances in Veterinary Science. 10:65–126.

10. Abd el-Rahim IH, Abd el-Hakim U, Hussein M. 1999. An epizootic of Rift Valley fever in Egypt in 1997. Rev Sci Tech. 18(3):741–8.

11. Davies FG, Martin V. Recognizing Rift Valley Fever. Rome: Food and Agriculture Organization of the United Nations, 2003.

12. Yedloutschnig RJ, Dardiri AH, Mebus CA, Walker JS. 1981. Abortion in vaccinated sheep and cattle after challenge with Rift Valley fever virus. Vet Rec. 109(17):383–4.

13. Adam I, Karsany MS. 2008. Case report: Rift valley fever with vertical transmission in a pregnant Sudanese woman. Journal of Medical Virology. 80(5):929-.

14. Arishi HM, Aqeel AY, Al Hazmi MM. 2006. Vertical transmission of fatal Rift Valley fever in a newborn. Annals of tropical paediatrics. 26(3):251–3.

15. Baudin M, Jumaa AM, Jomma HJE, Karsany MS, Bucht G, Naslund J, Ahlm C, Evander M, Mohamed N. 2016. Association of Rift Valley fever virus infection with miscarriage in Sudanese women: a cross-sectional study. Lancet Glob Health. 4(11):e864–e71.

16. Bird BH, Albarino CG, Nichol ST. 2007. Rift Valley fever virus lacking NSm proteins retains high virulence in vivo and may provide a model of human delayed onset neurologic disease. Virology. 362(1):10–5. PMCID: PMCPMID: 17412386.

17. Lane-Petter W. 1968. Cannibalism in rats and mice. Proc R Soc Med. 61(12):1295–6. PMCID: PMCPMC2211618.

18. Reynolds RD. 1981. Preventing maternal cannibalism in rats. Science. 213(4512):1146.

19. Bird BH, Bawiec DA, Ksiazek TG, Shoemaker TR, Nichol ST. 2007. Highly sensitive and broadly reactive quantitative reverse transcription-PCR assay for high-throughput detection of Rift Valley fever virus. J Clin Microbiol. 45(11):3506–13. PMCID: PMC2168471.

20. Platt DJ, Smith AM, Arora N, Diamond MS, Coyne CB, Miner JJ. 2018. Zika virus-related neurotropic flaviviruses infect human placental explants and cause fetal demise in mice. Sci Transl Med. 10(426).

21. Bales JM, Powell DS, Bethel LM, Reed DS, Hartman AL. 2012. Choice of inbred rat strain impacts lethality and disease course after respiratory infection with Rift Valley Fever Virus. Frontiers in cellular and infection microbiology. 2:105. PMCID: PMC3417668.

22. Peters CJ, Slone TW. 1981. Inbred rat strains mimic the disparate human response to Rift Valley fever virus infection. J Med Virol. 10(1):45–54.

23. Caroline AL, Kujawa MR, Oury TD, Reed DS, Hartman AL. 2015. Inflammatory Biomarkers Associated with Lethal Rift Valley Fever Encephalitis in the Lewis Rat Model. Front Microbiol. 6:1509. PMCID: PMCPMC4703790.

24. Soares MJ, Chakraborty D, Karim Rumi MA, Konno T, Renaud SJ. 2012. Rat placentation: an experimental model for investigating the hemochorial maternal-fetal interface. Placenta. 33(4):233–43. PMCID: PMCPMC3288880.

25. Segal L, Thacker K, Fochesato M, Giordano G, Garçon N, Destexhe E. 2017. Intramuscularly administered herpes zoster subunit vaccine has no effects on fertility, pre- and post-natal development in Sprague-Dawley rats. Reproductive Toxicology. 69:297–307.

26. Segal L, Wilby OK, Willoughby CR, Veenstra S, Deschamps M. 2011. Evaluation of the intramuscular administration of Cervarix™ vaccine on fertility, pre- and post-natal development in rats. Reproductive Toxicology. 31(1):111–20.

27. Ohkawara T, Katsuyama T, Ida-Eto M, Narita N, Narita M. 2015. Maternal viral infection during pregnancy impairs development of fetal serotonergic neurons. Brain and Development. 37(1):88–93.

28. Swanson AM, David AL. 2015. Animal models of fetal growth restriction: Considerations for translational medicine. Placenta. 36(6):623–30.

29. Pijnenborg R, Robertson WB, Brosens I, Dixon G. 1981. Review article: trophoblast invasion and the establishment of haemochorial placentation in man and laboratory animals. Placenta. 2(1):71–91.

30. World Health Organization. Rift Valley Fever fact sheet 2017. Available from: http://www.who.int/mediacentre/factsheets/fs207/en/.

31. Coetzer JA. 1982. The pathology of Rift Valley fever. II. Lesions occurring in field cases in adult cattle, calves and aborted foetuses. Onderstepoort J Vet Res. 49(1):11–7.

32. Szaba FM, Tighe M, Kummer LW, Lanzer KG, Ward JM, Lanthier P, Kim IJ, Kuki A, Blackman MA, Thomas SJ, Lin JS. 2018. Zika virus infection in immunocompetent pregnant mice causes fetal damage and placental pathology in the absence of fetal infection. PLoS pathogens. 14(4):e1006994. PMCID: PMCPMC5909921.

33. Suffin SC, Muck KB, Porter DD. 1977. Vesicular stomatitis virus causes abortion and neonatal death in ferrets. J Clin Microbiol. 6(4):437–8. PMCID: PMCPMC274789.

34. Arora N, Sadovsky Y, Dermody TS, Coyne CB. 2017. Microbial Vertical Transmission during Human Pregnancy. Cell host & microbe. 21(5):561–7.

35. Kourtis AP, Read JS, Jamieson DJ. 2014. Pregnancy and infection. The New England journal of medicine. 370(23):2211–8. PMCID: PMCPMC4459512.

36. Silasi M, Cardenas I, Kwon JY, Racicot K, Aldo P, Mor G. 2015. Viral Infections During Pregnancy. American Journal of Reproductive Immunology. 73(3):199–213.

37. Mosby LG, Rasmussen SA, Jamieson DJ. 2011. 2009 pandemic influenza A (H1N1) in pregnancy: a systematic review of the literature. American journal of obstetrics and gynecology. 205(1):10–8.

38. Neuzil KM, Reed GW, Mitchel EF, Simonsen L, Griffin MR. 1998. Impact of influenza on acute cardiopulmonary hospitalizations in pregnant women. Am J Epidemiol. 148(11):1094–102.

39. Price ME, Fisher-Hoch SP, Craven RB, McCormick JB. 1988. A prospective study of maternal and fetal outcome in acute Lassa fever infection during pregnancy. Bmj. 297(6648):584–7. PMCID: PMCPMC1834487.

40. Mupapa K, Mukundu W, Bwaka MA, Kipasa M, De Roo A, Kuvula K, Kibadi K, Massamba M, Ndaberey D, Colebunders R, Muyembe-Tamfum JJ. 1999. Ebola hemorrhagic fever and pregnancy. J Infect Dis. 179 Suppl 1:S11–2.

41. Niklasson B, Liljestrand J, Bergstrom S, Peters CJ. 1987. Rift Valley fever: a sero-epidemiological survey among pregnant women in Mozambique. Epidemiology and infection. 99(2):517–22. PMCID: PMCPMC2249273.

42. Gommet C, Billecocq A, Jouvion G, Hasan M, Zaverucha do Valle T, Guillemot L, Blanchet C, van Rooijen N, Montagutelli X, Bouloy M, Panthier JJ. 2011. Tissue tropism and target cells of NSs-deleted rift valley fever virus in live immunodeficient mice. PLoS neglected tropical diseases. 5(12):e1421. PMCID: PMC3232203.

43. Coetzer JA, Barnard BJ. 1977. Hydrops amnii in sheep associated with hydranencephaly and arthrogryposis with wesselsbron disease and rift valley fever viruses as aetiological agents. Onderstepoort J Vet Res. 44(2):119–26.

44. Crane J, Mundle W, Boucoiran I, Gagnon R, Bujold E, Basso M, Bos H, Brown R, Cooper S, Gouin K, McLeod NL, Menticoglou S, Pylypjuk C, Roggensack A, Sanderson F. 2014. Parvovirus B19 Infection in Pregnancy. Journal of Obstetrics and Gynaecology Canada. 36(12):1107–16.

45. Gao YL, Gao Z, He M, Liao P. 2018. Infection status of human parvovirus B19, cytomegalovirus and herpes simplex Virus-1/2 in women with first-trimester spontaneous abortions in Chongqing, China. Virology Journal. 15(1).

46. Sarno M, Sacramento GA, Khouri R, do Rosário MS, Costa F, Archanjo G, Santos LA, Nery N, Vasilakis N, Ko AI, de Almeida ARP. 2016. Zika Virus Infection and Stillbirths: A Case of Hydrops Fetalis, Hydranencephaly and Fetal Demise. PLoS neglected tropical diseases. 10(2).

47. Valentine G, Marquez L, Pammi M. 2016. Zika virus epidemic: an update. Expert Review of Anti-Infective Therapy. 14(12):1127–38.

48. Bayer A, Lennemann NJ, Ouyang Y, Bramley JC, Morosky S, Marques ET, Jr., Cherry S, Sadovsky Y, Coyne CB. 2016. Type III Interferons Produced by Human Placental Trophoblasts Confer Protection against Zika Virus Infection. Cell host & microbe. 19(5):705–12. PMCID: PMCPMC4866896.

49. Jagger BW, Miner JJ, Cao B, Arora N, Smith AM, Kovacs A, Mysorekar IU, Coyne CB, Diamond MS. 2017. Gestational Stage and IFN-lambda Signaling Regulate ZIKV Infection In Utero. Cell host & microbe. 22(3):366–76 e3. PMCID: PMCPMC5647680.

50. Delorme-Axford E, Donker RB, Mouillet JF, Chu T, Bayer A, Ouyang Y, Wang T, Stolz DB, Sarkar SN, Morelli AE, Sadovsky Y, Coyne CB. 2013. Human placental trophoblasts confer viral resistance to recipient cells. Proc Natl Acad Sci U S A. 110(29):12048–53. PMCID: PMCPMC3718097.

51. Corry J, Arora N, Good CA, Sadovsky Y, Coyne CB. 2017. Organotypic models of type III interferon-mediated protection from Zika virus infections at the maternal-fetal interface. Proc Natl Acad Sci U S A. 114(35):9433–8. PMCID: PMCPMC5584447.

52. Grigsby PL. 2016. Animal Models to Study Placental Development and Function throughout Normal and Dysfunctional Human Pregnancy. Semin Reprod Med. 34(1):11–6. PMCID: PMCPMC4799492.

53. Oreshkova N, van Keulen L, Kant J, Moormann RJ, Kortekaas J. 2013. A single vaccination with an improved nonspreading Rift Valley fever virus vaccine provides sterile immunity in lambs. PloS one. 8(10):e77461. PMCID: PMCPMC3805595.

54. Makoschey B, van Kilsdonk E, Hubers WR, Vrijenhoek MP, Smit M, Wichgers Schreur PJ, Kortekaas J, Moulin V. 2016. Rift Valley Fever Vaccine Virus Clone 13 Is Able to Cross the Ovine Placental Barrier Associated with Foetal Infections, Malformations, and Stillbirths. PLoS neglected tropical diseases. 10(3).

